# The genomic impact of mycoheterotrophy: targeted gene losses but extensive expression reprogramming

**DOI:** 10.1101/2020.06.26.173617

**Authors:** Jakalski Marcin, Minasiewicz Julita, Caius José, Michał May, Selosse Marc-André, Delannoy Etienne

## Abstract

Mycoheterotrophic plants have lost the ability to photosynthesize and they parasitize their associated fungus to get the mineral and organic nutrients they need. Despite involving radical changes in life history traits and ecological requirements, the transition from autotrophy to mycoheterotrophy occurred independently in almost all major lineages of land plants, but most often in *Orchidaceae*. Yet the molecular mechanisms underlying this shift are still poorly understood. The comparison of the transcriptomes of *Epipogium aphyllum* and *Neottia nidus-avis*, two mycoheterotrophic orchids, to other autotrophic and mycoheterotrophic orchids showed massive molecular function losses restricted to photosynthetic activities. In addition to these targeted losses, the analysis of their expression profiles showed that many orthologs had inverted root/shoot ratios compared to autotrophic species. Fatty acid and amino acid biosynthesis as well as primary cell wall metabolism were among the pathways most impacted by this expression reprogramming. Our study suggests that, while associated with function losses rather than metabolic innovations, the shift in nutritional mode from autotrophy to mycoheterotrophy remodeled the architecture of the plant metabolism.

## INTRODUCTION

More than 85% of vascular plants grow in association with soil fungi, forming a mycorrhizal symbiosis (Brundrett and Tedersoo, 2018; van der Heijden et al., 2015). Thanks to this symbiosis, plant growth and fitness are substantially improved by better mineral nutrition and increased resistance to biotic and abiotic stresses. In this mutualism, the fungal partner exchanges mineral nutrients (water, N, P, K…) against organic compounds from the photosynthesis of the plant (Rich et al., 2017). However, more than 500 plant species, called mycoheterotrophs (MH), have lost their ability to photosynthesize and entirely rely on their fungal partners to get both their mineral and organic nutrients, reversing the usual direction of the net carbon flow. The metabolic evolution associated with this switch, which is still poorly understood, has occurred in parallel at least 50 times in 17 plant families, including at least 30 independent transitions among *Orchidaceae* (Merckx et al., 2009; Těšitel et al., 2018; Merckx and Freudenstein, 2010). One of the characteristics that may make orchids more prone to evolutionary change of nutrition mode lies in their minute seeds devoid of nutritional reserves (Rasmussen, 1995). They fully depend on their mycorrhizal fungi for carbon supply at germination and early developmental stages (protocorms), which are thus always MH (Merckx, 2013; Dearnaley et al., 2016). Most orchid species later shift to autotrophy once photosynthesis becomes possible and establish more reciprocal exchanges with mycorrhizal fungi at adulthood. However, some species still recover carbon from the fungi at adulthood in addition to their photosynthesis (mixotrophy; Selosse and Roy, 2009), and from this nutrition some even evolved achlorophylly and mycoheterotrophy at adulthood (Dearnaley et al., 2016; Merckx et al., 2009). This versatile relation between orchids and their mycorrhizal partners provides an ideal framework to understand the metabolic evolution resulting in mycoheterotrophy (Lallemand et al., 2019; Suetsugu et al., 2017).

The impact of mycoheterotrophy on plant physiology can be analyzed through the changes in genomes of mycoheterotrophs (MH) compared to autotrophs (AT). As full mycoheterotrophy is associated with the loss of photosynthesis, sequencing of the plastid genome has been targeted first, thanks to next-generation methods (DePamphilis and Palmer, 1990; Delannoy et al., 2011; Bellot and Renner, 2016; Schelkunov et al., 2015). A common feature among MH plastid genomes is a strong reduction in size and gene content, especially with, as expected, a loss of all photosynthetic genes (Hadariová et al., 2018; Graham et al., 2017). However, the plastid genome contains only a tiny fraction of plant genes and the absence of genes from the plastid genome does not rule out the possibility that some of them were transferred into the nuclear genome, rather than lost (Bock, 2017).

Furthermore, in addition to photosynthesis, the transition to mycoheterotrophy can be expected to affect other metabolic processes, which cannot be assessed without the complete gene repertoire. Out of three published full genomes of achlorophyllous plants, two belong to obligate plant parasites (Vogel et al., 2018, Yoshida et al., 2019) and one to an east Asian mycoheterotrophic orchid (*Gastrodia elata* Blume; Yuan et al., 2018). When compared to photosynthetic orchids, the genome of *G. elata* is characterized by a reduction of the gene content, including the loss of most of the genes associated with photosynthesis, and the reduction of gene families involved in resistance to pathogens. At the same time, it shows an expansion of gene families putatively involved in the interaction with fungi (Yuan et al., 2018).

Despite the decrease in sequencing costs, the *de novo* characterization of a complete plant genome is still tedious and expensive, especially in the case of relatively large genomes of achlorophyllous orchids (from about 6 Gbases for *Corallorhiza trifida* Chatelain to about 16 Gbases for *Neottia nidus-avis* (L.) L.C.M. Rich; Pellicer and Leitch, 2020). Another approach for studying gene content is to analyze the transcriptome. RNA-seq focuses on the transcribed fraction of the genome, which includes the protein-coding genes. Transcriptomes of 5 mycoheterotrophic plants are currently available (Leebens-Mack et al., 2019; Schelkunov et al., 2018). The transcriptome of two orchids, *Epipogium aphyllum* Sw. and *E. roseum* (D. Don) Lindl., and the Ericaceae *Monotropa hypopitys* L. showed the loss of the photosynthetic genes (Schelkunov et al., 2018). Surprisingly, but in accordance with results from obligate parasitic plants (Wickett et al., 2011; Chen et al., 2020), the chlorophyll synthesis pathway was mostly conserved even if incomplete. However, transcriptome analysis only identifies the genes expressed in the tissue(s) under study, and since the previous studies of MH concentrated on the aerial part only, a fraction of the extant genes was likely missed. In addition, the missed genes include all the genes specifically expressed in the roots and mycorrhiza, which are fundamental to understanding of the mechanism of the interaction between an MH and its fungal partners. The switch to mycoheterotrophy not only results in gene losses, but also probably in neofunctionalizations and changes in the expression profiles of some retained genes, which are not captured in the analyses of gene repertoires.

In this study, we explored the transcriptome and gene expression profiles in the mycorrhiza, stems, and flowers of the MH orchids *Neottia nidus-avis* and *Epipogium aphyllum* (Figure 1). Both studied species are achlorophyllous and, like *G. elata*, belong to the *Epidendroideae* subfamily. Despite their rarity, they have a wide Eurasian range (Hulten and Fries, 1986) and, together with *G. elata*, they represent three independent evolutionary events of mycoheterotrophy (Merckx and Freudenstein, 2010). Their shoots with minute achlorophyllous scales support a few large flowers in *E. aphyllum* (Taylor and Roberts, 2011) and many small flowers in *N. nidus-avis* (Selosse, 2003). Considering their underground parts, *N. nidus-avis* displays a clump of short and thick mycorrhizal roots growing out of a short and thin rhizome, while *E. aphyllum* forms a fleshy, dichotomously branched and rootless rhizome that hosts the fungus (Roy et al., 2009). Thus, roots in *N. nidus-avis* and the rhizome of *E. aphyllum* are colonized by the mycorrhizal fungus.

**Figure 1:**
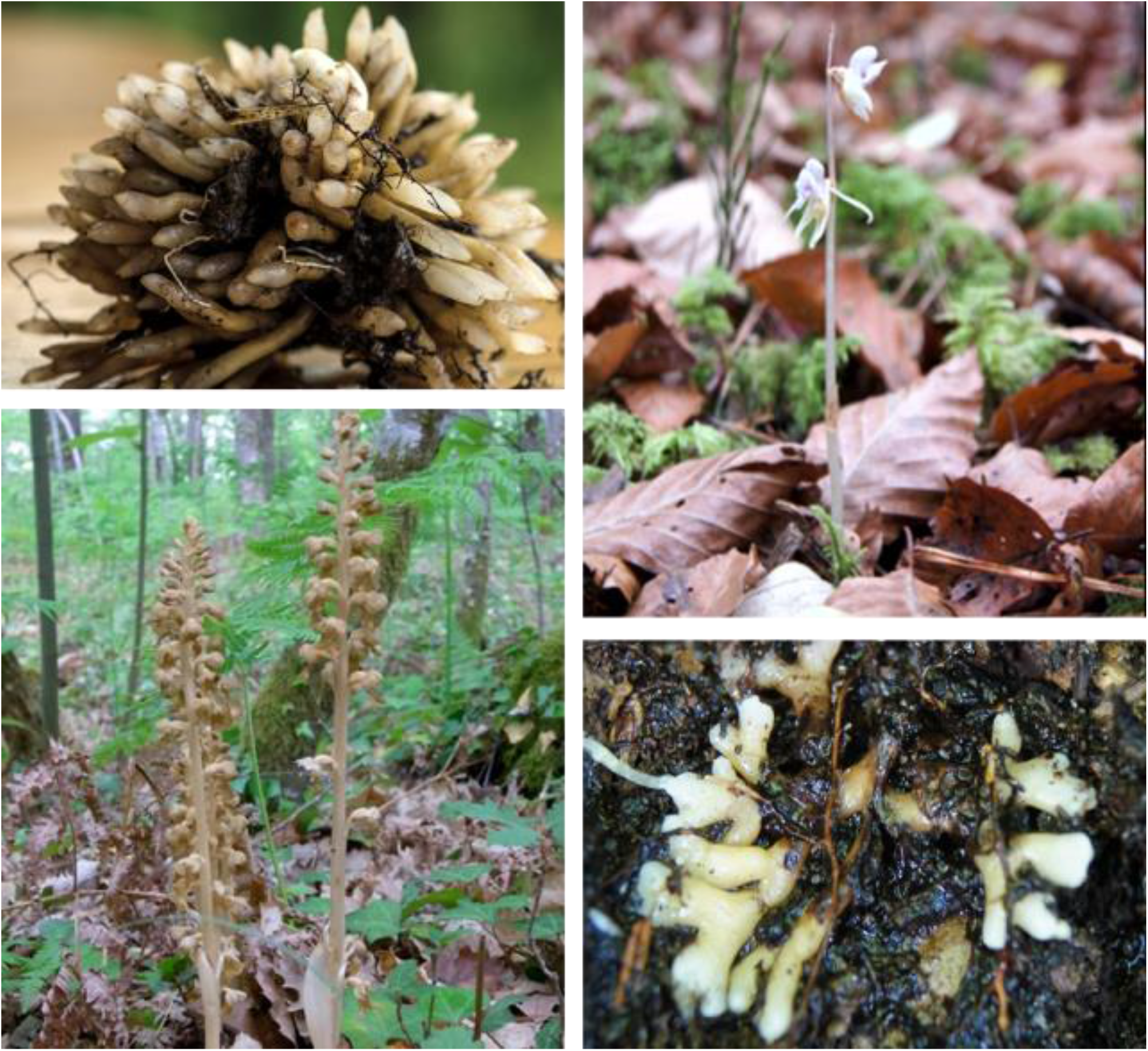
Morphology of *Neottia nidus-avis* and *Epipogium aphyllum*. Top left: roots of *N. nidus-avis*. Bottom left: inflorescence of *N. nidus-avis*. Top right: inflorescence of *E. phylum*. Bottom right: rhizome of *E. aphyllum*.

Using RNA-seq in flowers, stems, and mycorrhizal parts (roots or rhizomes) sampled in natural forest conditions, we identified expressed gene sets. In combination with the data from *G. elata*, we compared the gene sets of the three mycoheterotrophic orchids to that of three autotrophic orchid species, in order to highlight the gene losses and gains associated with the switch to mycoheterotrophy in orchids. We also identified the genes differentially expressed between the three investigated tissues. As no equivalent dataset (expression profiles per organ for the same individuals with biological replicates) is available for autotrophic orchids, we compared these profiles to expression in other autotrophic non-orchid plants. This comparison suggested that, in addition to gene losses, the switch to mycoheterotrophy induced extensive expression reprogramming.

## MATERIALS AND METHODS

### Sampling procedures

The specimens of *N. nidus-avis* and *E. aphyllum* were sampled in their natural habitats in southern Poland in 2017 at the peak of the species’ flowering season, at 10.00 am (Supplemental Table 1; these two species cannot be cultivated *ex situ*). Two plants per species were selected as biological replicates based on their size and healthy condition, keeping the parameters similar among the replicates. Each plant was excavated with surrounding soil. A fully open flower and the stem were cut off and processed (see below) right away, while the root system was first cleaned thoroughly by gentle scrubbing and rinsing in distilled water to remove all visible soil and foreign material. In-field processing consisted of slicing and dividing material into samples of 150 mg in weight before immediate preservation in liquid nitrogen to inhibit RNA degradation. The presence of mycorrhizal fungus was checked on thin cross-sections of colonized organs adjacent to the sampling and examined later under a light microscope.

### RNA extraction and purification

The samples collected *in situ* were transferred from liquid nitrogen to a −80°C freezer until the RNA extraction step. Flower samples were homogenized in liquid nitrogen using TissueLyser II (Qiagen) in 2 mL Eppendorf tubes containing ceramic beads. This method has proven ineffective in processing root, rhizome, and stem tissue samples due to their hardness when frozen. Manual grinding in mortars with liquid nitrogen had to be applied instead. Homogenized material was subjected to the NucleoZol (Macherey-Nagel, Dueren, Germany) reagent extraction process following the manufacturer’s protocol with the addition of polyvinylpolypyrrolidone (PVPP) during the grinding of root and rhizome. RNAs were further purified using Agencourt RNAClean XP (Beckman Coulter, Brea, CA, USA) magnetic beads following the manufacturer’s instructions.

Digestion with DNase Max (Qiagen, Hilden, Germany) was subsequently conducted to purify RNA isolates from remaining genomic DNA contamination.

Finally, RNA integrity and purity were assessed by Agilent BioAnalyzer 2100 survey using the Plant RNA Nano Chip (Agilent Technology, Santa Clara, CA, USA) and RNA concentration was measured by RiboGreen assay (Thermo Fisher Scientific, Waltham, MA, USA). Samples exhibiting high integrity (RIN > 7) were selected for sequencing.

### Sequencing

The RNA sequencing analyses of the isolated samples were performed at the Institute of Plant Sciences Paris-Saclay (IPS2, Saclay, France). First, the sequencing libraries were constructed using TruSeq Stranded Total RNA with the Ribo-Zero Plant kit (Illumina, San Diego, CA, USA), following the manufacturer’s instructions. Next, they were sequenced on a NextSeq500 (Illumina,

San Diego, CA, USA) platform in a paired-end mode with read length of 150 bases. The sequences obtained were quality-controlled and trimmed using the Trimmomatic software (version 0.36, parameters PE and ILLUMINACLIP:TruSeq3-PE.fa:2:30:10:2:true LEADING:21 TRAILING:21 MINLEN:30) (Bolger et al., 2014) and the residual ribosomal RNAs were filtered out with SortMeRNA (version 2.1, against the databases silva-bac-16s-id90, silva-bac-23s-id98, silva-euk-18s-id95 and silva-euk-28s-id98 with the parameters - e 1e-07 --paired_in).

### Transcriptome assembly

Due to the lack of reference genomes of the sampled plants, their transcriptomes were assembled *de novo* using the Trinity RNA-seq assembler (Haas et al., 2013) version v2.6.6 with all parameters at their default settings except --SS_lib_type RF. Taking into account the possible contamination of our samples collected *in situ*, which may vary between collected tissues, in order to avoid miss-assemblies and/or chimeric transcript generation, we assembled the transcriptomes of each collected plant tissue separately, but pooled both replicates. Subsequently, the assemblies for each species were merged and the redundancy of the isoforms was decreased with the tr2aacds pipeline from the EvidentialGene package (http://arthropods.eugenes.org/EvidentialGene/trassembly.html, version 2017.12.21; Gilbert, 2019). According to the pipeline description, we kept only those contigs that were classified as “main” or “noclass”, i.e. primary transcripts with alternates or with no alternates, respectively, to form the final unigene set.

### Identification of contaminating contigs

As our samples were collected in the field, the total RNAs contain transcripts from the microbiotas of our plants, especially abundant transcripts of the mycorrhizal fungi in underground organs, which means that the previous unigene set is contaminated with sequences not belonging to the species of interest.

To identify and filter out these contigs, the reduced transcriptome was searched with the blastx algorithm against the NCBI NR database using Diamond software (Buchfink et al., 2015). Local Diamond version 0.9.16 installation was run with the following set of parameters: --sensitive, --index-chunks 2, --block-size 20, --max-target-seqs 50, --no-self-hits, --evalue 0.001, --taxonmap prot.accession2taxid.gz, with the latest parameter specifying the taxonomic information obtained from the NCBI ftp pages (ftp://ftp.ncbi.nlm.nih.gov/pub/taxonomy/). Both the NCBI NR database and the taxonomy information were current as of December 2018. All contigs with best hits inside the *Streptophyta* clade of plants were considered as bona fide orchid contigs.

However, this analysis may miss genes highly conserved across kingdoms. Hence, we performed an orthology analysis against several orchid and monocotyledon species. The analysis included proteomes of *N. nidus-avis* and *E. aphyllum* generated here, as well published reference sets of *Brachypodium distachyon* (L.) P.Beauv.*, Zea mays* L.*, Oryza sativa*, and of the orchids *G. elata, Dendrobium catenatum* Lindley*, Apostasia shenzhenica* Z.J.Liu & L.J.Chen and *Phalaneopsis equestris* (Schauer) Rchb.f. (see Supplemental Table 2). We identified orthologous groups using the OrthoFinder software (version 2.2.7, default parameters, except -S diamond) (Emms and Kelly, 2019).

Contigs sharing the same orthogroup as any protein of these 7 species were considered as *bona fide* orchid contigs. For contigs with no hit at all we applied a further filtering criterion based on the expression pattern, i.e. we required such transcripts to be expressed in at least two out of our 6 samples. Expression analyses were performed with Seal from the BBTools package (https://jgi.doe.gov/data-and-tools/bbtools/, version 38.22).

### Identification of the fungal partners

The contig sets were searched for ITS sequences using ITSx software (version 1.1.2 (Bengtsson-Palme et al., 2013)) and the identified contigs were queried against the UNITE database (version 8.2, (Nilsson et al., 2019)).

### Annotations

Annotation of transcripts was performed with the Trinotate suite (version 3.1.1, https://trinotate.github.io/; Bryant et al., 2017). Trinotate was fed with results of several independent analyses. To annotate protein domains, hmmscan from the HMMER 3.1b2 package (Eddy, 2011) was run against the Pfam-A entries from the PFAM database (Finn et al., 2016). The UniProt/SwissProt protein database was searched with blastp (Altschul et al., 1997) from the blast+ 2.7.1 package to retrieve e.g. gene ontology (GO), KEGG (Kyoto Encyclopedia of Genes and Genomes), and eggNOG annotations. The presence of signal peptides was assessed with signalP (Petersen et al., 2011) software.

Additionally, the transcripts were assigned to KEGG orthologs and pathways using the KAAS server (Moriya et al., 2007) with BLAST and the BBH (bi-directional best hit) method. They were also assigned to the Mapman4 pathways using the Mercator4 v2.0 online tool (Schwacke et al., 2019).

In all the above analyses, transcripts were represented by either their nucleotide sequences derived directly from the assembly or by their amino acid sequences, as derived from the open reading frames (ORFs) determined by the tr2aacds pipeline. To avoid any technical bias when comparing species, the gene sets of all species were re-annotated with the same tools and parameters. The annotation of the orthogroups was derived from the annotations of their genes independently of the origin of these genes. If a term was present in more than 25% of its genes, the orthogroup was annotated with this term.

### Comparison of gene sets

The quality and completeness of the final transcriptomes (unigene sets) for *E. aphyllum* and *N. nidus-avis* were benchmarked with BUSCO v3.0.2 (Seppey et al., 2019) against the *Liliopsida*:odb10 plant-specific reference database and compared with the abovementioned species. We also compared the representation of the KEGG pathways and Mapman4 bins in each species. The unigene sets of *E. aphyllum* and *N. nidus-avis* were first completed with their plastid gene lists extracted from the NCBI accessions NC_026449.1 and NC_016471.1, respectively. We counted whether a KEGG ortholog or its Mapman equivalent was detected independently of the number of genes associated with it. Fisher’s exact test was performed to compare *E. aphyllum* and *N. nidus-avis* to *G. elata* and to compare these three mycoheterotrophic orchids to the three autotrophic orchids in each pathway or bin. Pathways or bins with an adjusted p-value (Bonferroni adjustment) below 0.05 were considered as differentially represented.

### Gene expression analyses

Sequencing read libraries were mapped separately to their corresponding final transcriptome (unigene set) using BBmap (https://jgi.doe.gov/data-and-tools/bbtools/). The software was run with the additional ‘rpkm’ parameter, which yields per-contig raw counts directly along the standard SAM/BAM output files. Next, a raw count matrix was generated for each species’ unigene set and fed into edgeR (Robinson et al., 2010) for differential expression testing by fitting a negative binomial generalized log-linear model (GLM) including a tissue factor and a replicate factor to the TMM-normalized read counts for each unigene. Unigenes detected in less than 3 of the 6 samples were considered as poorly expressed and filtered out from the analysis. We performed pairwise comparisons of tissues, i.e. flower vs. mycorrhiza (FL vs. MR), flower vs. stem (FL vs. ST), and mycorrhiza vs. stem (MR vs. ST). The distribution of the resulting p-values followed the quality criterion described by Rigaill et al. (2018). Genes with an adjusted p-value (FDR, Benjamini-Hochberg adjustment (1995)) below 0.05 were considered as differentially expressed.

Given the sets of up- and down-regulated genes for each species from pairwise tissue comparisons, we performed enrichment analysis for GO terms, KEGG and Mapman4 pathways using hypergeometric tests. Terms with an adjusted p-value (Bonferroni adjustment) below 0.05 were considered as enriched.

### Comparison of shoot/root expression profiles between autotrophs and mycoheterotrophs

As no equivalent dataset is available for autotrophic orchids, we used datasets from *Z. mays* and *B. distachyon* as autotrophic species for comparison. We focused on the root and stem tissues using roots and internodes as the corresponding tissues for autotrophic monocotyledons. Expression values for *Z. mays* were extracted from the SRA project PRJNA217053. The samples SRR957475 and SRR957476 correspond to internodes, SRR957460 and SRR957461 to roots. Expression values for *B. distachyon* were extracted from the SRA project PRJNA419776. The samples SRR6322422 and SRR6322429 correspond to internodes, SRR6322386 and SRR6322417 to roots. Counts were calculated after mapping of the reads to their corresponding reference transcriptome (Zea_mays.B73_RefGen_v4.cdna.all.fa and Brachypodium_distachyon.Brachypodium_distachyon_v3.0.cdna.all.fa) using BBmap with the same parameters as previously.

To allow a direct comparison between species, we used the expression values per orthogroup using the sum of counts of the orthogroup members in a given sample and species, an approach analogous to that of McWhite et al. (2020). Any orthogroup expression of which was not detected in at least one sample of all four species was filtered out from further analysis. As an orthogroup can group different numbers of genes from each species, the absolute counts cannot be directly compared. However, as the shoot and root samples are paired, it is possible to compare the root/shoot ratios. After normalization with the TMM method (Robinson et al., 2010) to correct the library size effect, the counts were transformed with the vst method of the coseq package v1.2 (Rau and Maugis-Rabusseau, 2018). The log2 root/shoot ratios calculated from the transformed counts were analyzed using the lmFit contrasts.fit and eBayes functions of the limma package v3.34.9 (Smyth, 2004). In our model, the log2 ratio was expressed as a linear combination of a species effect and the p-values corresponding to the difference between the average of MH and the average of AT were calculated. The distribution of the resulting p-values followed the quality criterion described by Rigaill et al. (2018). The Benjamini-Hochberg correction was used to control false discovery rate. We considered orthogroups with an adjusted p-value < 0.05 as having a different shoot/root ratio between AT and MH. Enrichment analyses were performed as described previously with orthogroups being annotated with terms representing at least 25% of their genes.

### Data availability

The reads are available at the NCBI database under Bioproject PRJNA633477. The GFF file and annotation of the unigene sets for *E. aphyllum* and *N. nidus-avis* as well as the raw count matrices are available at https://doi.org/10.15454/HR9KUX.

## RESULTS

### Sequencing, de novo assembly and functional annotation

To characterize the variability of RNA resulting from tissue- and species-specific features among the two studied mycoheterotrophic orchids sampled *in natura*, we performed Illumina short-read sequencing and *de novo* assembly of their transcriptomes. In total, 12 cDNA libraries from flowers, stems, and mycorrhizal roots (two replicates per tissue and species) were sequenced which resulted in 304,280,061 reads for *N. nidus-avis* and 178,486,849 reads for *E. aphyllum*. After assembly and filtering of the probable contaminating transcripts, the final set of transcripts comprised 44,451 and 38,488 sequences for *N. nidus-avis* and *E. aphyllum*, respectively (Table 1). As expected, the fraction of contaminating contigs was much higher in the mycorrhizal samples (roots in *N. nidus-avis* and rhizome in *E. aphyllum*), and indeed almost all the contaminating transcripts were most probably of fungal origin (Supplemental Tables 3 and 4). Thanks to the presence of a few contigs corresponding to ITS, the main fungal partners could be identified as *Inocybe cervicolor* (Pers.) Quél. and *Hebeloma incarnatulum* A.H. Sm. for *E. aphyllum* and *Sebacina epigaea* (Berk. & Broome) Bourdot & Galzin for *N. nidus-avis* as expected (McKendrick et al., 2002; Selosse et al., 2002; Roy et al., 2009)

**Table 1:** Statistics of the final assemblies.

|  | *Neottia nidus-avis* | *Epipogium aphyllum* |
| --- | --- | --- |
| Number of transcripts | 43 451 | 38 488 |
| Number of “genes” | 39 731 | 36 275 |
| Median / mean transcript length (bp) | 675 / 993 | 555 / 920 |
| Shortest / longest transcript (bp) | 201 / 13 537 | 201 / 15 415 |
| Transcripts over 1k / 10k bp length | 15 391 / 10 | 11 980 / 9 |
| Transcript n50 (bp) | 1 506 | 1 480 |
| GC % | 45.56 | 44.26 |
| Total assembled bases | 43 160 528 | 35 412 792 |

The generated transcripts of the two studied species were functionally annotated, mainly on the basis of homology to known reference sequences. The annotations included, among others, information on encoded protein domains (Pfam), Gene Ontology (GO) classification, KEGG Orthology (KO), metabolic pathway membership and Mapman4 mapping. Roughly 46% and 50% of the transcripts could be attributed to any annotation category in *N. nidus-avis* and *E. aphyllum*, respectively (Supplemental Table 5, Supplemental Data 1).

The completeness of the generated transcriptomes was assessed through several analyses. The BUSCO (Benchmarking Universal Single-Copy Orthologs) analysis showed 78.5% and 71% of completeness for *N. nidus-avis* and *E. aphyllum*, respectively, which is comparable to or higher than that for the *G. elata* genome (73.1%) and much higher than that for the *E. aphyllum* transcriptome generated by Schelkunov et al. (2018) (53.4%; Supplemental Table 6). We also checked the mapping rate of the RNA-seq reads on these transcriptomes (Supplemental Table 7). It was higher than 94%, except for mycorrhizal samples as expected because of the presence of the mycorrhizal fungi. Finally, we looked for the orthologs of the plant KEGG pathways and of the Mapman4 bins and compared them to the *G. elata* gene content (Supplemental Data 2). An ortholog was counted if at least one transcript or gene was associated with it. Out of the 140 tested KEGG pathways representing 15150 potential orthologs, none were differentially represented between our transcriptomes and *G. elata*. Identically, none of the 1196 tested Mapman4 bins representing 4966 potential orthologs were differentially represented. Even when relaxing the stringency of the test (raw p-value <0.05), no bin or pathway suggesting missed orthologs in our transcriptomes compared to *G. elata* was statistically significant. Taken together, these results strongly support that our transcriptomes were complete and that any missing ortholog from our transcriptomes was lost by the corresponding species. It also suggests that the gene repertoires of *E. aphyllum*, *N. nidus-avis* and *G. elata* are similar.

### Impact of mycoheterotrophy on the gene repertoire

Considering three mycoheterotrophic orchids of independent evolutionary origin, we can study the impact of mycoheterotrophy on the gene sets. A comparison with the genomes of *P. equestris*, *D. catenatum* and *A. shenzhenica*, three autotrophic orchids, using the KEGG and Mapman4 pathways described previously (Table 2, Supplemental Data 2), showed that the switch to mycoheterotrophy results in the loss of orthologs exclusively associated with pathways directly related to photosynthesis. Even when relaxing the stringency of the test (raw p-value <0.05), there is no indication that the switch to mycoheterotrophy could be associated with any gain (Supplemental Data 2). It should be noted that none of the genes lost from their plastid genomes were found in the transcriptomes of *E. aphyllum* and *N. nidus-avis*.

**Table 2:**
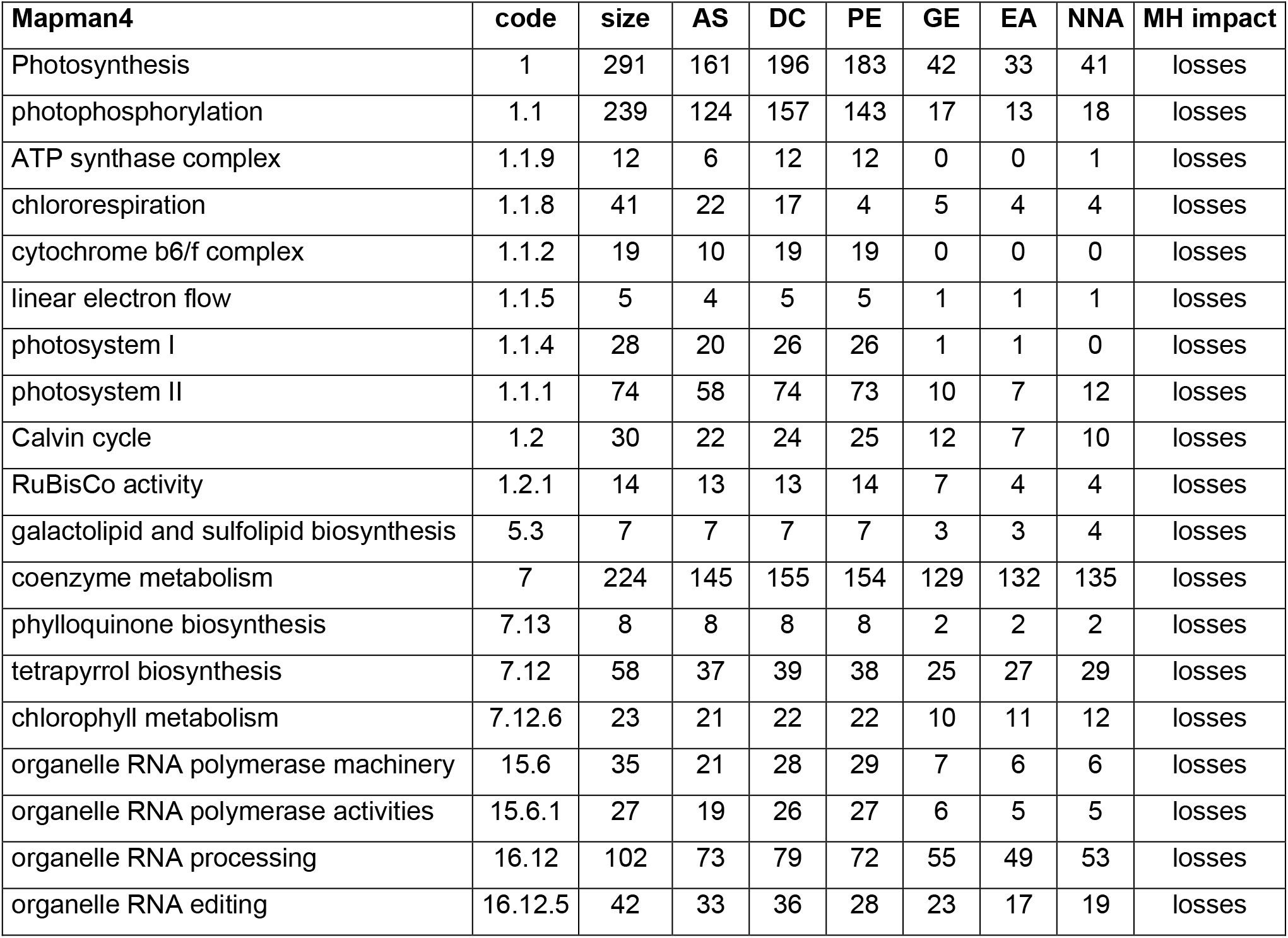
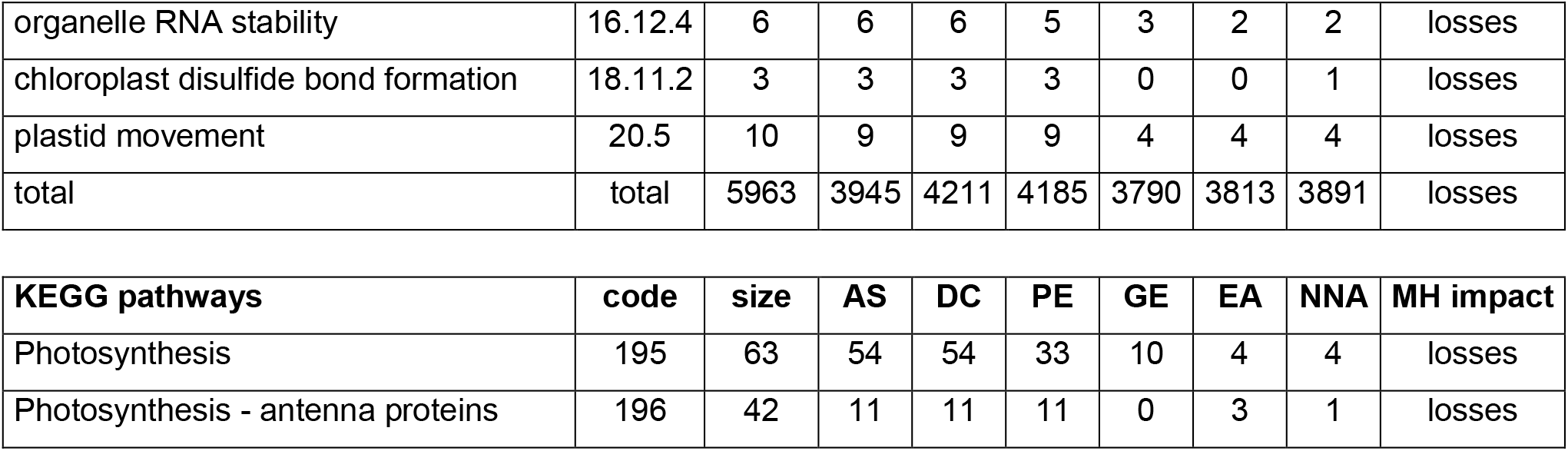
Gene content: pathways impacted by the switch to mycoheterotrophy. Code: code of the Mapman4 bin or KEGG pathway. Size: ortholog content of the bin or pathway. AS: *A. shenzhenica*. DC: *D. catenatum*. PE: *P. equestris*. GE: *G. elata*. EA: *E. aphyllum*. NNA: *N. nidus-avis*. MH impact: impact of the switch to mycoheterotrophy.

All the orthologs required for photosystems were lost, but the losses in the chlorophyll metabolism pathway were almost exclusively restricted to chlorophyll degradation and interconversion. As seen before (Schelkunov et al., 2018; Wickett et al., 2011), the chlorophyll synthesis pathway was mostly conserved but incomplete in *G. elata* and *E. aphyllum* (Figure 2A). On the other hand, *N. nidus-avis* expressed the full extent of genes required for the biosynthesis of chlorophyll as well as some chlorophyll a/b binding proteins (Light-Harvesting-Complex A3 (LHCA3), LHCB1, LHCB2, Stress-enhanced protein 1 (SEP1), SEP3, SEP5 and early light-induced protein (ELIP) genes). Similarly, the three MH species were missing the *lycE* and *lut5* genes required for the synthesis of lutein, but possessed a complete biosynthesis pathway to violaxanthin (Figure 2B). It should be noted that no gene coding for a violaxanthin de-epoxidase was found in any of the 3 MH species.

**Figure 2:**
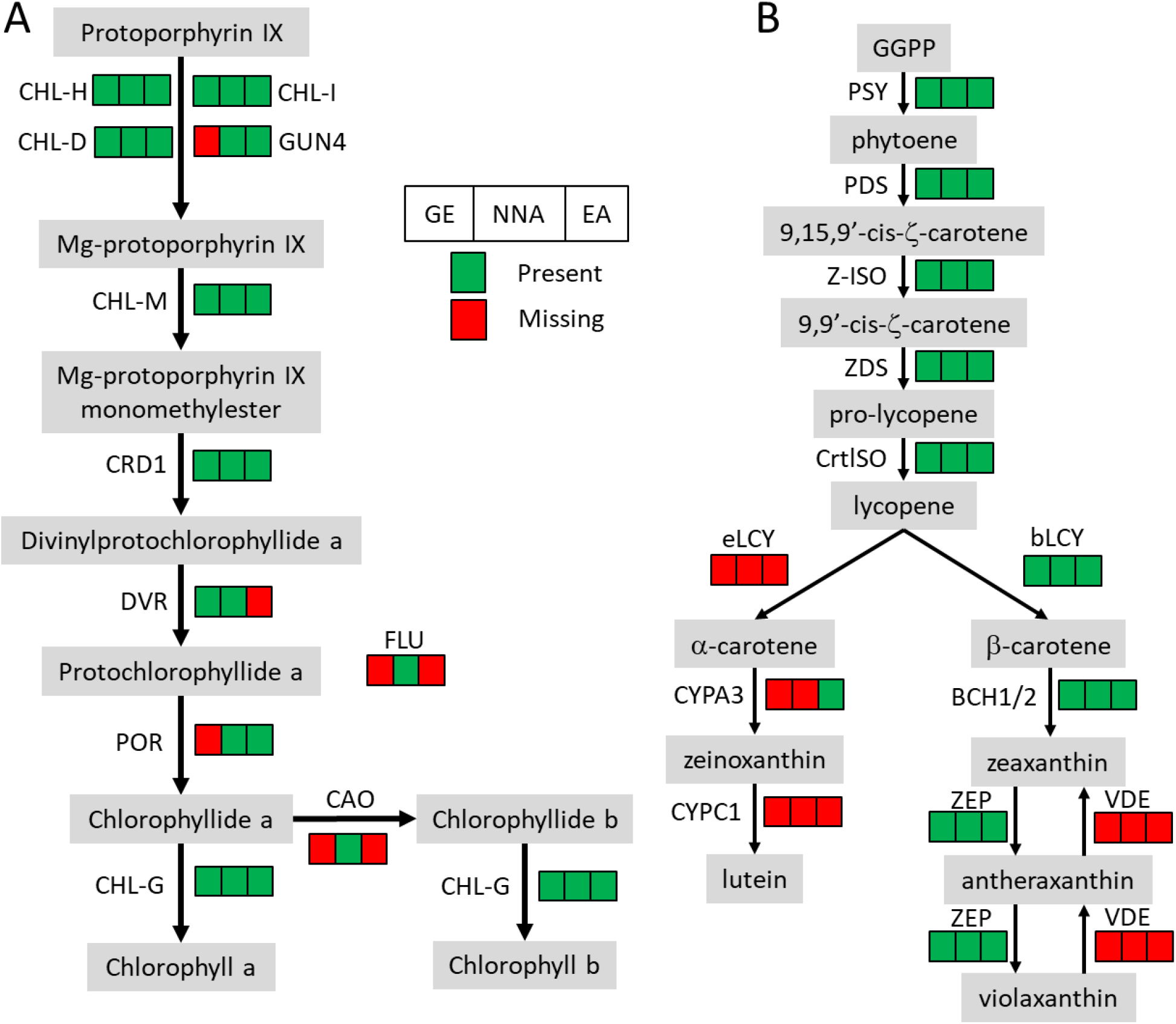
Pigment synthesis pathways in mycoheterotrophic orchids. GE: *G. elata*. NNA: *N. nidus-avis*. EA: E*. aphyllum*. A: Chlorophyll biosynthesis. CHL-D, CHL-H, CHL-I, GUN4: magnesium chelatase. CHL-M: Mg-protoporphyrin IX O-methyltransferase. CRD1: Mg-protoporphyrin IX monomethylester cyclase. DVR: divinyl chlorophyllide-a 8-vinyl-reductase. POR: protochlorophyllide oxidoreductase. FLU: glutamyl-tRNA reductase regulator. CAO: chlorophyllide a oxygenase. CHL-G: chlorophyll synthase. B: Carotenoid biosynthesis. PSY: Phytoene synthase. PDS: phytoene desaturase. Z-ISO: ζ-carotene isomerase. ZDS: ζ-carotene desaturase. CrtISO: carotenoid isomerase. eLCY: lycopene ε-cyclase. bLCY: lycopene β-cyclase. BCH1/2: β-ring carotene hydroxylase. ZEP: zeaxanthin epoxidase. VDE: violaxanthin de-epoxidase. CYPA3: α-carotene β-ring hydroxylase. CYPC1: carotenoid ε-hydroxylase.

Even when relaxing the stringency of the analysis, only pathways associated with plastid or photosynthesis were identified as lost, suggesting that the switch to mycoheterotrophy selectively impacted activities associated with photosynthesis. So, the switch to mycoheterotrophy significantly impacted pathways associated with photosynthesis, but targeted losses can be observed as well.

Using known pathways mainly based on autotrophic plants allows identification of gene losses linked to the switch to mycoheterotrophy, but misses potential new pathways or genes specifically associated with mycoheterotrophy. To overcome this problem, we performed an orthology analysis including the coding genes of the 6 previous orchid species plus *Z. mays*, *B. distachyon* and *O. sativa* (Supplemental Data 3 and Supplemental Tables 8 and 9). Out of the 18259 orthogroups identified, only 38 contained exclusively genes from all 3 MH orchid species. Twenty-two of these orthogroups contained only unannotated genes and the 16 remaining did not show specific annotations (Supplemental Data 4). These results suggest that the switch to mycoheterotrophy in orchids does not involve new pathways or functions.

### A transcriptome analysis highlights the organ-specific functions of mycoheterotrophic orchids

The pairwise comparisons of the transcriptome profiles of flower, stem, and mycorrhizal root of *E. aphyllum* and *N. nidus-avis* identified the genes differentially expressed between these organs as well as organ-specific genes (Supplemental Data 5). We identified 18817 and 12331 differentially expressed genes as well as 6351 and 4520 organ-specific genes in *N. nidus-avis* and *E. aphyllum*, respectively (Table 3). The highest numbers of differentially expressed genes were observed between underground and aerial organs. Similarly, most organ-specific genes were identified in the mycorrhizal root.

**Table 3:** Summary of differential gene expression analyses among the sampled tissues.

|  | *Neottia nidus-avis* | *Epipogium aphyllum* |
| --- | --- | --- |
| stem vs. flower | 9109 / 4644 down, 4465 up | 5315 / 2123 down, 3192 up |
| mycorrhiza vs. flower | 13701 / 6465 down, 7236 up | 7596 / 3430 down, 4166 up |
| mycorrhiza vs. stem | 11360 / 4866 down, 6494 up | 7849 / 3955 down, 3894 up |
| Flower-specific | 55 | 297 |
| Stem-specific | 508 | 175 |
| Mycorrhiza-specific | 5788 | 4048 |
| Total | 25168 (57.92%) | 16851 (43.78%) |

To elucidate which functions are served by the differentially expressed and organ-specific genes, Gene Ontology, Mapman and KEGG enrichment analyses were performed (Supplemental Data 6). While very few enrichments were found in the organ-specific genes, the differentially expressed genes showed that numerous metabolic functions were differentially activated in the 3 organs, following a strikingly similar pattern in *N. nidus-avis* and *E. aphyllum*. Figure 3 summarizes the Mapman and KEGG enrichment analyses, which are fully supported by the GO enrichment analyses. The metabolic functions are indicated where their activity appears to be peaking. The aerial parts shared high amino acid and fatty acid syntheses as well as high primary cell wall metabolism. They also activated light signaling pathways. The flowers specifically showed high cell division and phenolic activities. In *N. nidus-avis*, the activity of the chlorophyll synthesis pathway in association with other plastid activities was detected mostly in the flowers. At the other end of the plant, the mycorrhizal roots mostly showed an increased activity of pathways related to pathogen and symbiont interactions, as well as of the transportome (e.g. ABC transporters and solute carriers) and degradation capacities (proteasome and glycosaminoglycan and trehalose degradations).

**Figure 3:**
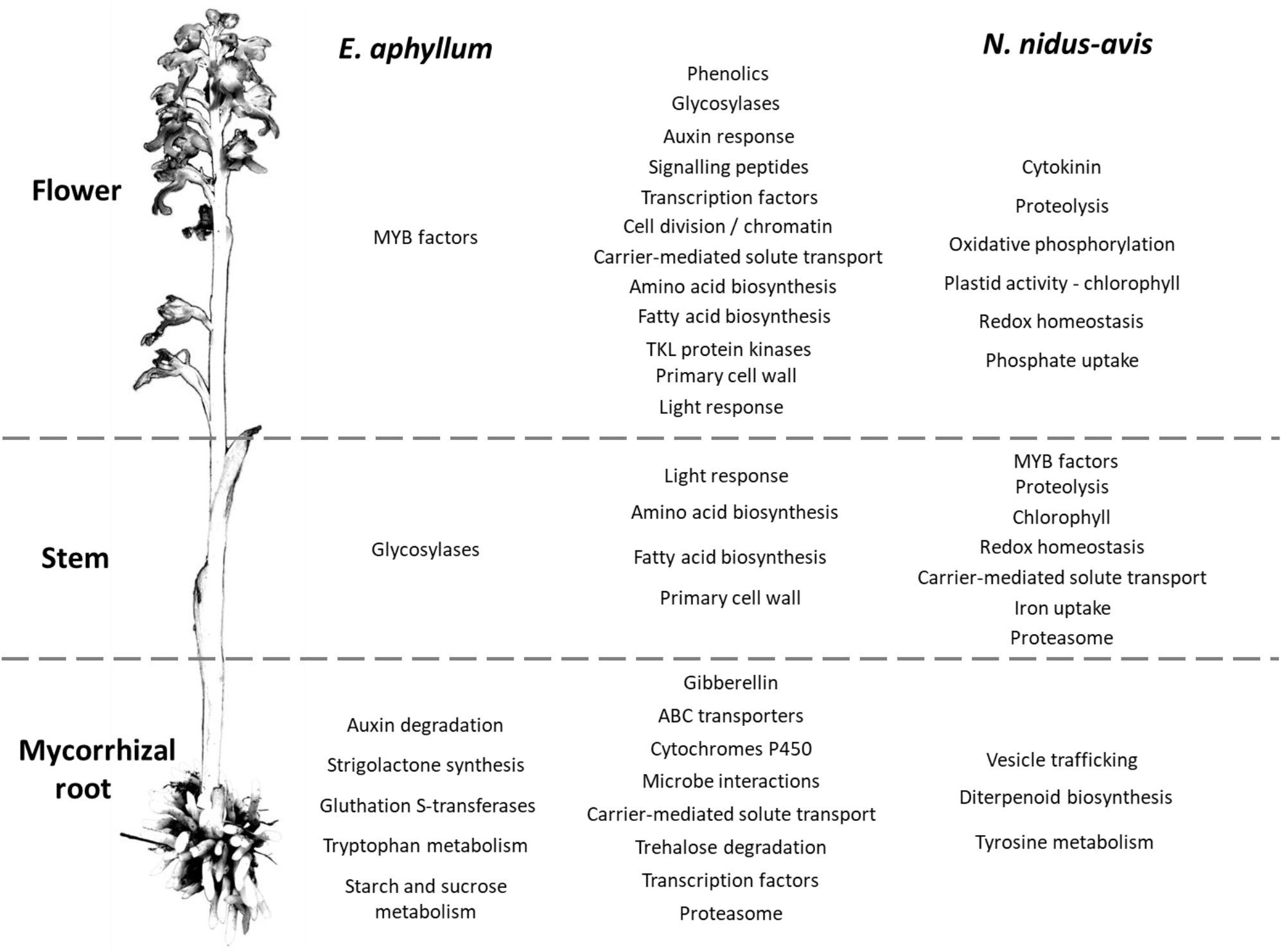
Pathways differentially expressed between organs in *E. aphyllum* and *N. nidus-avis*. Summary of the enrichment analysis of the transcriptomic expression profiles (Supplemental Data 5). A pathway is indicated in the organ(s) where its activity peaks. The common changes are shown in the central column while the changes specific to *E. aphyllum* (resp. *N. nidus-avis*) are shown in the left (resp. right) column. The terms are mostly derived from the Mapman4 and KEGG pathways.

### Comparison of expression profiles in roots and stem of mycoheterotrophic and autotrophic species

To understand the consequences of mycoheterotrophy for the expression profiles, it is necessary to compare our mycoheterotrophic orchids to AT species from a transcriptomic point of view. As no equivalent transcriptomic dataset is publicly available for autotrophic orchids, we used datasets from two other monocots, *B. distachyon* and maize. However, rather than analyzing them as previously to compare the enrichment analyses for our species, we directly compared the 4 species using the orthogroup expression levels. As the number and length of the genes in each orthogroup can differ from one species to another, we compared the root/stem ratios of expression in MH and AT. We analyzed only the 8620 (out of 18259) orthogroups detected in the roots or stem of all four species. This filter removes most of the orthogroups associated with photosynthesis, but these pathways are an obvious difference between the two trophic types.

While 2378 and 3617 orthogroups were differentially expressed between root and stem in AT and MH, respectively, 3359 orthogroups showed a significantly different root to stem ratio between the 2 trophic types, including 2536 with inverted ratios (Supplemental Data 7).

The pathway enrichment analysis of the differentially expressed orthogroups in MH (Supplemental Data 8) showed results similar to the transcriptomic analysis of *E. aphyllum* and *N. nidus-avis* genes, supporting the fact that the analysis of orthogroup expression is biologically relevant. Even with the exclusion of most photosynthesis-related orthogroups, we observed a different root to stem ratio between AT and MH for almost 39% of the analyzed orthogroups and even an inversion of this ratio for 30% of the orthogroups. Figure 4 summarizes the results of the pathway enrichment analysis of these orthogroups. It is particularly noteworthy that the fatty acid, amino acid, and primary cell wall metabolisms, which are high in the stem of MH, are actually higher in the root of AT. In addition, this analysis highlighted that glycosidases and the secondary metabolism seemed higher in the stem of MH, but lower in the stem of AT, while the opposite was true for RNA metabolism and DNA damage response. Some pathways (solute transport, symbiosis, trehalose degradation and cytochrome P450) were more expressed in the roots than in the stems for both AT and MH, but differed between AT and MH, suggesting that the species of the two trophic types either induced these pathways to different levels or used different orthologs.

**Figure 4:**
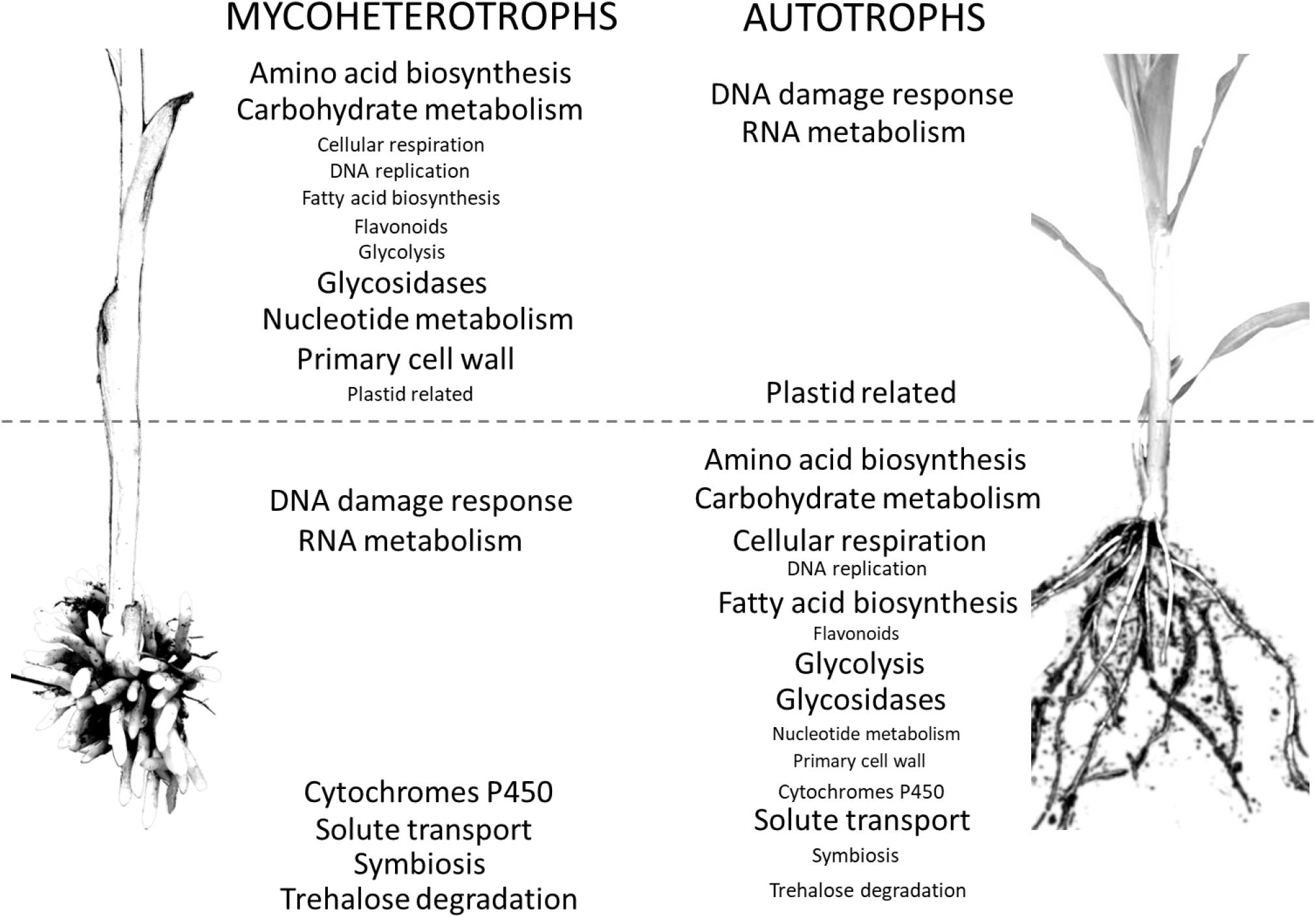
Comparison of distribution of pathways between the organs of mycoheterotrophic orchids and autotrophic non-orchid plants. Pathways enriched in the orthogroups showing a significantly different root/stem expression ratio between mycoheterotrophic species (*N. nidus-avis* and *E. aphyllum*) and autotrophic species (*B. distachyon* and *Z. mays*). The pathways are indicated in the organ where their expression is highest. The pathways shown with a large font are also differentially expressed between root and stem.

The latter can be illustrated for the “solute transport” pathway. The 192 orthogroups showing a different root/shoot ratio between AT and MH (out of 392 orthogroups belonging to the solute transport pathway) are distributed in most transporter families, and in each family there are orthologs showing different behavior in MH and AT (Supplemental Data 7). All this shows that the consequences of mycoheterotrophy extend well beyond photosynthesis and the gene losses observed previously. Mycoheterotrophy remodeled a large fraction of gene expression and metabolism.

## DISCUSSION

Mycoheterotrophic species that rely entirely on their fungal partners for their nutrition (Merckx et al., 2009) reverse the usual mycorrhizal exchange, where fungi receive plant carbon. The switch to mycoheterotrophy, which involves the loss of photosynthesis, a hallmark of plants which is central to their metabolism, occurred at least 30 times in the orchid family and remains poorly understood at the molecular level. Independent events in a similar phylogenetic context can be used to study the impact of this metabolic evolution. We studied this question in orchids through transcriptome analysis in organs of *N. nidus-avis* and *E. aphyllum*, two MH orchids representing independent occurrences of mycoheterotrophy within Epidendroideae. We compared their gene content to that of other orchids, but also their expression profile to autotrophic orchids and non-orchid species.

### No genetic innovation, but gene loss

Using the RNA-seq data from rhizome/root, stem and flowers of *E. aphyllum* and *N. nidus-avis*, we identified gene sets that are probably almost complete, based on their comparison with the genome of *G. elata* (Supplemental Data 2). When comparing the molecular functions encoded in these 3 MH orchids with those of AT orchids, the switch to mycoheterotrophy entailed a global reduction of molecular functions, as previously demonstrated for *G. elata* (Supplemental Data 2; Yuan et al., 2018). In addition, we could not detect any major gain of function associated with mycoheterotrophy. Obviously, it is difficult to identify potentially unknown functions and our transcriptome analysis must have missed some genes, including those specifically expressed during germination. However, all orchids are mycoheterotrophic during germination (Dearnaley et al., 2016; Merckx, 2013), so mycoheterotrophy is not specific to MH species. Moreover, this behavior of autotrophic orchids during germination indirectly indicates that they have all the genes and metabolic pathways required to obtain nutrients through mycoheterotrophic nutrition, showing that major gains/innovation are not essential for the transition to mycoheterotrophy. However, when looking for orthologs present in our 3 MH species, but not in the other 6 AT species, we found only a handful of MH-specific orthologs. It is highly unlikely that they are the key genes required for the switch to mycoheterotrophy, although more extensive sampling of MH and AT species may verify this possibility.

In addition to a general reduction of gene content, Yuan et al. (2018) showed that some gene families, mostly associated with interactions with fungi, expanded in the *G. elata* genome. Our transcriptome assemblies include large numbers of contigs putatively coding for enzymes such as mannose-specific lectins or β-glucosidases, indicating the possible expansion of some gene families in *E. aphyllum* and *N. nidus-avis*. However, using transcriptome assemblies and despite a step of redundancy reduction in our analysis, it is difficult to count the number of genes precisely because it is impossible to distinguish between 2 isoforms and 2 copies of a gene. Only high-quality assemblies of the large genome of these species (2×16.96 Gb for *N. nidus-avis;* Vesely et al., 2012) can confirm the expansion of some gene families.

### Pigments and secondary metabolism: compensatory protection and camouflage?

The losses observed in MH species reflect the evolution of their plastomes, with massive gene loss restricted to photosynthetic pathways and functions. Genes retained in the plastid genomes have non-photosynthetic functions. By extension to the nuclear genome, the orthologs lost in MH species are probably exclusively associated with photosynthesis, while the orthologs conserved in MH species probably have non-photosynthetic functions. Comparison of the gene content of MH and AT species should provide interesting information for the functional analysis of genes even in model plants, as shown by two examples below.

The loss of photosynthesis resulted in gene losses in several pigment synthesis pathways (Table 2). In *N. nidus-avis*, Pfeifhofer (1989) detected high amounts of zeaxanthin but no lutein. In the three MH species, the genes coding for the enzymatic activities of the carotenoid pathway required for the synthesis of zeaxanthin, but not lutein, are conserved (Figure 2). Lutein is associated with the dissipation of excess energy from the photosystems and zeaxanthin is part of the xanthophyll cycle, which has the same function (Niyogi et al., 1997). However, the loss of violaxanthin de-epoxidase shows loss of the xanthophyll cycle in these species. The fact that zeaxanthin is also a precursor of abscisic acid can explain the conservation of a functional synthesis pathway. The switch to mycoheterotrophy trimmed the multifunctional carotenoid synthesis pathway to keep only the enzymes required for its non-photosynthetic functions.

Although MH species are expected to lose the chlorophyll synthesis pathway, it is nonetheless conserved, even if incomplete, in *E. aphyllum* and *G. elata* (Figure 2). Such a conservation was already observed in holoparasitic plants (Wickett et al., 2011), and suggests that chlorophylls or their intermediates could have a non-photosynthetic function, which is still not clear (Ankele et al., 2007). *N. nidus-avis* differs from the two other species by a complete and functional chlorophyll synthesis pathway. Its activity, in association with other plastid activities, was detected in *N. nidus-avis*, mostly in the flowers (Figure 3). This is consistent with the detection of chlorophyll a and b in the inflorescence (Pfeifhofer, 1989). Menke and Schmid (1976) reported a cyclic photophosphorylation in the flower of *N. nidus-avis*, but this report is incompatible with the absence of most plastid and nuclear genes coding for photosystem I and cytochrome b6/f and deserves further study.

Because of the photo-toxicity of free chlorophyll and their precursors (Rebeiz et al., 1984), the accumulation of chlorophyll requires a photo-protection mechanism. Flowers of *N. nidus-avis* are not green, but they turn green upon heating (Supplemental Figure 1), suggesting that the chlorophyll is stored in a heat-labile complex, which may limit toxicity. When compared with *G. elata* and *E. aphyllum*, the activity of the chlorophyll synthesis pathway in *N. nidus-avis* is associated with the presence of several SEP and ELIP genes. The SEP1 and ELIP Arabidopsis orthologs are induced in response to high light and are believed to bind chlorophyll (Heddad, 2000; Adamska et al., 1999; Rossini et al., 2006), but their exact molecular functions are unknown. Their conservation in *N. nidus-avis*, but not in E*. aphyllum* or *G. elata*, suggests that they may indeed bind chlorophyll to inactivate its ability to capture light.

Another, non-exclusive possible explanation for conservation of a functional chlorophyll synthesis pathway and the accumulation of zeaxanthin to high levels in *N. nidus-avis* (Pfeifhofer, 1989) may be camouflage. By blending the plants in the surrounding leaf litter, the dull colors of MH species protect them against herbivory (Klooster et al., 2009).

In any case, we show that the switch to mycoheterotrophy is mostly dominated by function losses, and does not require major, massive metabolic innovations. In mixotrophic species (an evolutionary transition from autotrophy to mycoheterotrophy; Selosse and Roy, 2009), a metabolomic and transcriptomic analysis showed that their response to the loss of photosynthesis was similar to the response of AT species to achlorophylly (Lallemand et al., 2019). This suggests that the ability of achlorophyllous variants of otherwise green mixotrophic species to sustain an almost normal growth without photosynthesis is mostly based on the plasticity of plant metabolism. Furthermore, mycoheterotrophy is not a rare event (it has occurred > 50 times in 17 plant families; Merckx et al., 2009; Těšitel et al., 2018), suggesting that it entails function losses rather than complex gene gains.

### An upside-down metabolic architecture

Being at the core of plant metabolism, the loss of photosynthesis in normally green plants severely impacts their metabolism (Lallemand et al., 2019; Aluru et al., 2009; Abadie et al., 2016). The switch to mycoheterotrophy remodels the genome and we analyzed MH physiology through gene expression in different organs (Figure 3). As expected, many genes were differentially expressed, reflecting a partition of metabolic functions between the organs. The flowers showed a higher activity of cell division, primary cell wall and signaling pathways, which can be attributed to floral development. Similarly, higher phenolic compound synthesis can be associated with pollinator attraction involving flower pigmentation and production of fragrant phenolics (Jakubska-Busse et al., 2014). Conversely, the underground organs showed a higher activity of pathways likely involved in the interaction with their fungal partners (microbe interactions, proteasome, transporters).

Although *N. nidus-avis* and *E. aphyllum* showed similar pathway enrichments, especially in the aerial organs, there were some idiosyncrasies that may result from different phylogenetic backgrounds, as well as different fungal partners. The peak of tryptophan, starch and sucrose metabolism observed in the rhizome of *E. aphyllum* as opposed to a peak of tyrosine metabolism in the roots of *N. nidus-avis* can provide clues to the specificities of the nutrient fluxes in these two pairs of partners.

Comparing symbiotic and asymbiotic protocorms of *Serapias vomeracea*, Fochi et al. (2017) highlighted the importance of organic N metabolism and especially lysine histidine transporters (LST) in the interaction with the fungal partner. In our analysis, several LST genes were differentially expressed between the organs for both *N. nidus-avis* and *E. aphyllum*, but some were induced in flowers while others were more transcribed in stems or mycorrhizal parts (Supplemental Data 7). In a similar analysis in *G. elata*, the upregulation of clathrin genes in symbiotic protocorms suggested the involvement of exocytosis (Zeng et al. 2017). Our analysis showed no signal specific to N metabolism or exocytosis. The different conditions considered in these studies can explain the discrepancies, but they may also illustrate some evolutionary tinkering occurring in different mycorrhiza.

Comparison of mycoheterotrophs’ expression profiles to similar datasets in AT *B. distachyon* and maize provides additional evidence of the impact of mycoheterotrophy on plant metabolism. Its interpretation is limited by factors such as different phylogenetic backgrounds, possibly different growth conditions (incl. absence of mycorrhizal fungi), or the restriction to orthogroups detected in the four species. Yet almost 40% of the analyzed orthogroups had a significantly different root/stem ratio between MH and AT species, and 30% of the orthogroups, from numerous pathways, showed inverted root/shoot ratios, suggesting that MH metabolism was somehow upside-down. This inversion of the metabolism architecture coincided with the inversion of the source/sink relationship: in MH, underground organs are sources, while they are a sink in AT. The sink organs were associated with a higher activity of several major metabolic pathways (carbohydrate and nucleotide metabolism, amino acid and fatty acid biosynthesis, glycolysis, and respiration). In association with a higher DNA replication and primary cell wall activity (which involves glycosidases) and a higher expression of auxin transporters, sink organs likely experience stronger growth than their source counterparts. Mycoheterotrophic roots and rhizomes are generally short, thick and compact to minimize accidental loss of a part of a source organ and nutrient transfer effort (Imhof et al., 2013), stems are ephemeral (<2 months) but fast growing (4 cm/day in *E. aphyllum*, J. Minasiewicz pers. observations) sexual organs without nutritional functions. Conversely, fibrous roots of grasses have high growth rate as nutrient uptake depends largely on the root length (Fitter, 2002), while aerial internodes have much slower growth, or even stop growing.

Even with different growth habits, some pathways showed similar overall root/shoot ratios in AT and MH. Plastid-related pathways (chlorophyll synthesis, plastid translation) are more active in shoots than roots, while symbiosis and trehalose degradation are more active in roots than shoots. Trehalose is almost absent from vascular plants, where its 6-phosphaste precursor is an important growth regulator (Lunn et al., 2014). However, it is an abundant storage carbohydrate in mycorrhizal fungi and it has been suggested that it is transferred to host MH orchids to be cleaved into glucose (Müller and Dulieu, 1998). The comparison between leaves of achlorophyllous mutants (with MH nutrition) and green individuals in mixotrophic orchids showed an upregulation of trehalase, but also of trehalose-6-P phosphatases (TPP) and trehalose-6-P synthase (TPS; Lallemand et al. 2019). Similarly, the MH species showed a higher root/shoot ratio of trehalase and TPP (but not TPS) compared to AT, which supports the hypothesis that trehalose is transferred from mycorrhizal fungi to MH plants. Many other nutrients are exchanged at this interface and our analysis suggests numerous differences between AT and MH: close to half of the orthogroups involved in solute transport showed different root to stem ratios between AT and MH. Some SWEET transporters were induced in the mycorrhiza of achlorophyllous MH mutants of the mixotrophic orchid *E. helleborine* (Suetsugu et al., 2017) and in the protocorms of *Serapias* (Perotto et al., 2014). The three SWEET orthogroups in our analysis behaved differently between AT and MH, but showed contrasted differences, indicating that AT and MH both used SWEET transporters but different orthologs in roots and stem. Similarly, 13 out of the 15 ABCG transporter orthogroups or 10 out of the 13 NRT1/PTR transporter orthogroups showed contrasted differences between AT and MH. The same could be observed for all transporter families (Supplemental Data 7): AT and MH use different orthologs for the transport of solutes in stem and roots, demonstrating extensive expression reprogramming. These differences are probably associated with changes in the fluxes of nutrients in AT and MH, including in mycorrhizas.

This is a central question in the study of MH. However, the specificity of transporters can vary even within a family. For example, transporters of the NRT1/PTR family were identified as nitrate transporters, but some transport other molecules (Corratgé-Faillie and Lacombe, 2017). To understand the changes of nutrient fluxes associated with this reprogramming of transporter expression would require a detailed analysis of each orthogroup (assuming that the substrate specificity is the same for all transporters within an orthogroup). However, this analysis would not replace direct measurement of these fluxes with labeling experiments.

## CONCLUSIONS

The shift to mycoheterotrophy induces contrasting changes in the genome of MH plants. From the analysis of the gene repertoires, we were not able to identify new functions associated with mycoheterotrophy, and large losses seemed restricted to genes only involved in photosynthetic functions. This suggested that no metabolic innovation is required for mycoheterotrophy. However, the transcriptome analysis showed extensive changes in numerous pathways, probably associated with changes in the plant lifecycle and in the interaction with fungal partners induced by mycoheterotrophy. This reprogramming illustrates the versatility of plant metabolism and can be considered as a metabolic innovation by itself. It also explains why, since becoming MH is based more on reprogramming and gene loss than on genetic innovation, the shift to MH nutrition has occurred more than 50 times in plant evolution.

## Supplemental Data

**Supplemental Data 1**: Distribution of GO terms in the 3 mycoheterotrophic orchids.

**Supplemental Data 2**: Comparison of orthologue numbers in Mapman and KEGG pathways for the 3 mycoheterotrophic orchids and 3 autotrophic orchids.

**Supplemental Data 3**: Output of the Orthofinder analysis.

**Supplemental Data 4**: Composition and annotation of the mycoheterotroph-specific orthogroups.

**Supplemental Data 5**: Differential expression analysis of *N. nidus-avis* and *E. aphyllum* organs.

**Supplemental Data 6**: Mapman, KEGG and GO enrichment analysis of *N. nidus-avis* and *E. aphyllum* expression.

**Supplemental Data 7**: Differential analysis of the root/shoot expression ratios.

**Supplemental Data 8**: Mapman, KEGG and GO enrichment analysis of the expression ratios.

**Supplemental Table 1**: Details of sampling location and dates for the studied orchids.

**Supplemental Table 2**: Genomic datasets used in this study.

**Supplemental Table 3**: Comparison of the intermediate and final assemblies generated.

**Supplemental Table 4**: Composition of contamination sources among sampled tissues.

**Supplemental Table 5:** Annotation statistics of the generated transcriptome assemblies.

**Supplemental Table 6**: Summary statistics of the BUSCO analysis of completeness for the generated transcriptomes in comparison to the *E. aphyllum* transcriptome from Schelkunov et al. (2018) and another mycoheterotrophic orchid *G. elata* with a sequenced genome.

**Supplemental Table 7**: Statistics of per-tissue read mapping to the intermediate and final assemblies.

**Supplemental Table 8:** Per-species statistics among the generated orthologous groups.

**Supplemental Table 9**: Species overlaps among orthologous groups.

## Acknowledgements

This work was financially supported by grants from the National Science Center, Poland (project No: 2015/18/A/NZ8/00149). The IPS2 benefits from the support of Saclay Plant Sciences-SPS (ANR-17-EUR-0007). We thank Emilia Krawczyk for the photos of *E. aphyllum*.

## Author Contributions

MAS and ED designed the study, MAS supervised the project; ED, MM and MJ analyzed the data. ED, JM and MJ wrote the manuscript. JC generated the RNA-seq data. JM, MJ and MAS collected the samples.

## REFERENCES

Abadie, C., Lamothe-Sibold, M., Gilard, F., and Tcherkez, G. (2016). Isotopic evidence for nitrogen exchange between autotrophic and heterotrophic tissues in variegated leaves. Funct. Plant Biol. 43: 298.

Adamska, I., Roobol-Bóza, M., Lindahl, M., and Andersson, B. (1999). Isolation of pigment-binding early light-inducible proteins from pea. Eur. J. Biochem. 260: 453–460.

Altschul, S.F., Madden, T.L., Schäffer, A.A., Zhang, J., Zhang, Z., Miller, W., and Lipman, D.J. (1997). Gapped BLAST and PSI-BLAST: a new generation of protein database search programs. Nucleic Acids Res. 25: 3389–402.

Aluru, M.R., Zola, J., Foudree, A., and Rodermel, S.R. (2009). Chloroplast photooxidation-induced transcriptome reprogramming in Arabidopsis immutans white leaf sectors 1[w][OA]. Plant Physiol. 150: 904–923.

Ankele, E., Kindgren, P., Pesquet, E., and Strand, A. (2007). *In vivo* visualization of Mg-protoporphyrin IX, a coordinator of photosynthetic gene expression in the nucleus and the chloroplast. Plant Cell 19: 1964–1979.

Bellot, S. and Renner, S.S. (2016). The Plastomes of Two Species in the Endoparasite Genus *Pilostyles (Apodanthaceae)* Each Retain Just Five or Six Possibly Functional Genes. Genome Biol Evol 8: 189–201.

Bengtsson-Palme, J. et al. (2013). Improved software detection and extraction of ITS1 and ITS2 from ribosomal ITS sequences of fungi and other eukaryotes for analysis of environmental sequencing data. Methods Ecol. Evol. 4: n/a-n/a.

Benjamini, Y. and Hochberg, Y. (1995). Controlling The False Discovery Rate - A Practical And Powerful Approach To Multiple Testing. J. R. Stat. Soc. B 57: 289–300.

Bock, R. (2017). Witnessing Genome Evolution: Experimental Reconstruction of Endosymbiotic and Horizontal Gene Transfer. Annu. Rev. Genet. 51: 1–22.

Bolger, A.M., Lohse, M., and Usadel, B. (2014). Trimmomatic: a flexible trimmer for Illumina sequence data. Bioinformatics 30: 2114–2120.

Brundrett, M.C. and Tedersoo, L. (2018). Evolutionary history of mycorrhizal symbioses and global host plant diversity. New Phytol. 220: 1108–1115.

Bryant, D.M. et al. (2017). A Tissue-Mapped Axolotl *De Novo* Transcriptome Enables Identification of Limb Regeneration Factors. Cell Rep. 18: 762–776.

Buchfink, B., Xie, C., and Huson, D.H. (2015). Fast and sensitive protein alignment using DIAMOND. Nat. Methods 12: 59–60.

Chen, J., Yu, R., Dai, J., Liu, Y., and Zhou, R. (2020). The loss of photosynthesis pathway and genomic locations of the lost plastid genes in a holoparasitic plant *Aeginetia indica*. BMC Plant Biol. 20: 199.

Corratgé-Faillie, C. and Lacombe, B. (2017). Substrate (un)specificity of Arabidopsis NRT1/PTR FAMILY (NPF) proteins. J. Exp. Bot. 68: 3107–3113.

Dearnaley, J., Perotto, S., and Selosse, M.A. (2016). Structure and development of orchid mycorrhizas. Mol. Mycorrhizal Symbiosis: 63–86.

Delannoy, E., Fujii, S., Colas Des Francs-Small, C., Brundrett, M., and Small, I. (2011). Rampant Gene loss in the underground orchid *Rhizanthella gardneri* highlights evolutionary constraints on plastid genomes. Mol. Biol. Evol. 28.

DePamphilis, C.W. and Palmer, J.D. (1990). Loss of photosynthetic and chlororespiratory genes from the plastid genome of a parasitic flowering plant. Nature 348: 337–339.

Eddy, S.R. (2011). Accelerated Profile HMM Searches. PLoS Comput. Biol. 7: e1002195.

Emms, D.M. and Kelly, S. (2019). OrthoFinder: Phylogenetic orthology inference for comparative genomics. Genome Biol. 20.

Finn, R.D. et al. (2016). The Pfam protein families database: towards a more sustainable future. Nucleic Acids Res. 44: D279–85.

Fitter, A. (2002). Characteristics and Functions of Root Systems. In Plant roots - the hidden half, Y. Waisel, A. Eshel, and U. Kafkafi, eds (Marcel Dekker, Inc.), pp. 21–50.

Fochi, V., Chitarra, W., Kohler, A., Voyron, S., Singan, V.R., Lindquist, E.A., Barry, K.W., Girlanda, M., Grigoriev, I. V., Martin, F., Balestrini, R., and Perotto, S. (2017). Fungal and plant gene expression in the *Tulasnella calospora–Serapias vomeracea* symbiosis provides clues about nitrogen pathways in orchid mycorrhizas. New Phytol. 213: 365–379.

Gilbert, D.G. (2019). Longest protein, longest transcript or most expression, for accurate gene reconstruction of transcriptomes? bioRxiv: 1–27.

Graham, S.W., Lam, V.K.Y., and Merckx, V.S.F.T. (2017). Plastomes on the edge: the evolutionary breakdown of mycoheterotroph plastid genomes. New Phytol. 214: 48–55.

Haas, B.J. et al. (2013). *De novo* transcript sequence reconstruction from RNA-seq using the Trinity platform for reference generation and analysis. Nat. Protoc. 8: 1494–512.

Hadariová, L., Vesteg, M., Hampl, V., and Krajčovič, J. (2018). Reductive evolution of chloroplasts in non-photosynthetic plants, algae and protists. Curr. Genet. 64: 365–387.

Heddad, M. (2000). Light stress-regulated two-helix proteins in *Arabidopsis thaliana* related to the chlorophyll a/b-binding gene family. Proc. Natl. Acad. Sci. 97: 3741–3746.

van der Heijden, M.G.A., Martin, F.M., Selosse, M.A., and Sanders, I.R. (2015). Mycorrhizal ecology and evolution: The past, the present, and the future. New Phytol. 205: 1406–1423.

Hulten, E. and Fries, M. (1986). Atlas of North European Vascular Plant, North of the Tropic of Cancer.

Imhof, S., Massicotte, H.B., Melville, L.H., and Peterson, R.L. (2013). Subterranean morphology and mycorrhizal structures. In Mycoheterotrophy: The Biology of Plants Living on Fungi (Springer New York), pp. 157–214.

Jakubska-Busse, A., Jasicka-Misiak, I., Poliwoda, A., Świeczkowska, E., and Kafarski, P. (2014). The chemical composition of the floral extract of *Epipogium aphyllum* Sw. (Orchidaceae): A clue for their pollination biology. Arch. Biol. Sci. 66: 989–998.

Klooster, M.R., Clark, D.L., and Culley, T.M. (2009). Cryptic bracts facilitate herbivore avoidance in the mycoheterotrophic plant *monotropsis odorata (ericaceae).* Am. J. Bot. 96: 2197–2205.

Lallemand, F., Martin-Magniette, M.-L., Gilard, F., Gakière, B., Launay-Avon, A., Delannoy, É., and Selosse, M.-A. (2019). *In situ* transcriptomic and metabolomic study of the loss of photosynthesis in the leaves of mixotrophic plants exploiting fungi. Plant J. 98.

Leebens-Mack, J.H. et al. (2019). One thousand plant transcriptomes and the phylogenomics of green plants. Nature 574: 679–685.

Lunn, J.E., Delorge, I., Figueroa, C.M., Van Dijck, P., and Stitt, M. (2014). Trehalose metabolism in plants. Plant J. 79: 544–567.

Mckendrick, S.L., Leake, J.R., Taylor, D.L., and Read, D.J. (2002). Symbiotic germination and development of the myco-heterotrophic orchid *Neottia nidus-avis* in nature and its requirement for locally distributed *Sebacina* spp. New Phytol. 154: 233–247.

McWhite, C.D. et al. (2020). A Pan-plant Protein Complex Map Reveals Deep Conservation and Novel Assemblies. Cell 181: 460–474.e14.

Menke, W. and Schmid, G.H. (1976). Cyclic Photophosphorylation in the Mykotrophic Orchid *Neottia nidus-avis*. Plant Physiol. 57: 716–719.

Merckx, V., Bidartondo, M.I., and Hynson, N.A. (2009). Myco-heterotrophy: when fungi host plants. Ann. Bot. 104: 1255–1261.

Merckx, V. and Freudenstein, J. V. (2010). Evolution of mycoheterotrophy in plants: A phylogenetic perspective. New Phytol. 185: 605–609.

Merckx, V.S.F.T. (2013). Mycoheterotrophy: The Biology of Plants Living on Fungi.

Moriya, Y., Itoh, M., Okuda, S., Yoshizawa, A.C., and Kanehisa, M. (2007). KAAS: an automatic genome annotation and pathway reconstruction server. Nucleic Acids Res. 35: W182–W185.

Müller, J. and Dulieu, H. (1998). Enhanced growth of non-photosynthesizing tobacco mutants in the presence of a mycorrhizal inoculum. J. Exp. Bot. 49: 707–711.

Nilsson, R. et al. (2019). The UNITE database for molecular identification of fungi: handling dark taxa and parallel taxonomic classifications. Nucleic Acids Res. 47: D259–D264.

Niyogi, K.K., Björkman, O., and Grossman, A.R. (1997). The roles of specific xanthophylls in photoprotection. Proc. Natl. Acad. Sci. U. S. A. 94: 14162–14167.

Pellicer, J. and Leitch, I.J. (2020). The Plant DNA C-values database (release 7.1): an updated online repository of plant genome size data for comparative studies. New Phytol. 226: 301–305.

Perotto, S., Rodda, M., Benetti, A., Sillo, F., Ercole, E., Rodda, M., Girlanda, M., Murat, C., and Balestrini, R. (2014). Gene expression in mycorrhizal orchid protocorms suggests a friendly plant-fungus relationship. Planta 239: 1337–1349.

Petersen, T.N., Brunak, S., von Heijne, G., and Nielsen, H. (2011). SignalP 4.0: discriminating signal peptides from transmembrane regions. Nat. Methods 8: 785–6.

Pfeifhofer, H.W. (1989). Evidence for Chlorophyll b and Lack of Lutein in *Neottia nidus-avis* Plastids. Biochem. und Physiol. der Pflanz. 184: 55–61.

Rasmussen, H.N. (1995). Terrestrial orchids : from seed to mycotrophic plant (Cambridge University Press).

Rau, A. and Maugis-Rabusseau, C. (2018). Transformation and model choice for RNA-seq co-expression analysis. Brief. Bioinform. 19: 425–436.

Rebeiz, C.A., Montazer-Zouhoor, A., Hopen, H.J., and Wu, S.M. (1984). Photodynamic herbicides: 1. Concept and phenomenology. Enzyme Microb. Technol. 6: 390–396.

Rich, M.K., Nouri, E., Courty, P.E., and Reinhardt, D. (2017). Diet of Arbuscular Mycorrhizal Fungi: Bread and Butter? Trends Plant Sci. 22: 652–660.

Rigaill, G. et al. (2018). Synthetic data sets for the identification of key ingredients for RNA-seq differential analysis. Brief. Bioinform. 19.

Robinson, M.D., McCarthy, D.J., and Smyth, G.K. (2010). edgeR: a Bioconductor package for differential expression analysis of digital gene expression data. Bioinformatics 26: 139–140.

Rossini, S., Casazza, A.P., Engelmann, E.C.M., Havaux, M., Jennings, R.C., and Soave, C. (2006). Suppression of both ELIP1 and ELIP2 in Arabidopsis does not affect tolerance to photoinhibition and photooxidative stress. Plant Physiol. 141: 1264–1273.

Roy, M., Yagame, T., Yamato, M., Koji, I., Heinz, C., Faccio, A., Bonfante, P., and Selosse, M.-A. (2009). Ectomycorrhizal *Inocybe* species associate with the mycoheterotrophic orchid *Epipogium aphyllum* but not its asexual propagules. Ann. Bot. 104: 595–610.

Schelkunov, M.I., Penin, A.A., and Logacheva, M.D. (2018). RNA-seq highlights parallel and contrasting patterns in the evolution of the nuclear genome of fully mycoheterotrophic plants. BMC Genomics 19.

Schelkunov, M.I., Shtratnikova, V., Nuraliev, M., Selosse, M., Penin, A.A., and Logacheva, M.D. (2015). Exploring the limits for reduction of plastid genomes: a case study of the mycoheterotrophic orchids *Epipogium aphyllum* and *Epipogium roseum*. Genome Biol Evol 7: 1179–1191.

Schwacke, R., Ponce-Soto, G.Y., Krause, K., Bolger, A.M., Arsova, B., Hallab, A., Gruden, K., Stitt, M., Bolger, M.E., and Usadel, B. (2019). MapMan4: A Refined Protein Classification and Annotation Framework Applicable to Multi-Omics Data Analysis. Mol. Plant 12: 879–892.

Selosse, M.-A. and Roy, M. (2009). Green plants that feed on fungi: facts and questions about mixotrophy. Trends Plant Sci. 14: 64–70.

Selosse, M. (2003). La Néottie, une“mangeuse” d’arbres. L’Orchidophile 155: 21–31.

Selosse, M.A., Weiβ, M., Jany, J.L., and Tillier, A. (2002). Communities and populations of sebacinoid basidiomycetes associated with the achlorophyllous orchid *Neottia nidus-avis* (L.) L.C.M. Rich. and neighbouring tree ectomycorrhizae. Mol. Ecol. 11: 1831–1844.

Seppey, M., Manni, M., and Zdobnov, E.M. (2019). BUSCO: Assessing genome assembly and annotation completeness. In Methods in Molecular Biology (Humana Press Inc.), pp. 227–245.

Smyth, G.K. (2004). Linear models and empirical bayes methods for assessing differential expression in microarray experiments. Stat. Appl. Genet. Mol. Biol. 3.

Suetsugu, K., Yamato, M., Miura, C., Yamaguchi, K., Takahashi, K., Ida, Y., Shigenobu, S., and Kaminaka, H. (2017). Comparison of green and albino individuals of the partially mycoheterotrophic orchid *Epipactis helleborine* on molecular identities of mycorrhizal fungi, nutritional modes and gene expression in mycorrhizal roots. Mol. Ecol. 26: 1652–1669.

Taylor, L. and Roberts, D.L. (2011). Biological Flora of the British Isles: *Epipogium aphyllum* Sw. J. Ecol. 99: 878–890.

Těšitel, J., Těšitelová, T., Minasiewicz, J., and Selosse, M.A. (2018). Mixotrophy in Land Plants: Why To Stay Green? Trends Plant Sci. 23: 656–659.

Vesely, P., Bures, P., Smarda, P., and Pavlicek, T. (2012). Genome size and DNA base composition of geophytes: the mirror of phenology and ecology? Ann. Bot. 109: 65–75.

Vogel, A. et al. (2018). Footprints of parasitism in the genome of the parasitic flowering plant *Cuscuta campestris*. Nat. Commun. 9: 2515.

Wickett, N.J., Honaas, L.A., Wafula, E.K., Das, M., Huang, K., Wu, B., Landherr, L., Timko, M.P., Yoder, J., Westwood, J.H., and Depamphilis, C.W. (2011). Transcriptomes of the parasitic plant family *Orobanchaceae* reveal surprising conservation of chlorophyll synthesis. Curr. Biol. 21: 2098–2104.

Yoshida, S. et al. (2019). Genome Sequence of *Striga asiatica* Provides Insight into the Evolution of Plant Parasitism. Curr. Biol. 29: 3041–3052.e4.

Yuan, Y. et al. (2018). The *Gastrodia elata* genome provides insights into plant adaptation to heterotrophy. Nat. Commun. 9.

Zeng, X., Li, Y., Ling, H., Liu, S., Liu, M., Chen, J., and Guo, S. (2017). Transcriptomic analyses reveal clathrin-mediated endocytosis involved in symbiotic seed germination of *Gastrodia elata*. Bot. Stud. 58.

